# Ovarian Tumor Mitochondria Exhibit Abnormal Phenotypes and Blunted Associations with Biobehavioral Factors

**DOI:** 10.1101/2021.02.26.432917

**Authors:** Snehal Bindra, Marlon A. McGill, Marina K. Triplett, Anisha Tyagi, Premal H. Thaker, Laila Dahmoush, Michael J. Goodheart, R. Todd Ogden, Edward Owusu-Ansah, Kalpita Karan, Steve Cole, Anil K. Sood, Susan K. Lutgendorf, Martin Picard

## Abstract

Tumor cells exhibit mitochondrial alterations and are also influenced by biobehavioral processes, but the intersection of biobehavioral factors and tumor mitochondria remains unexplored. Here we examined multiple biochemical and molecular markers of mitochondrial content and function in benign and cancerous ovarian tissue in parallel with exploratory analyses of biobehavioral factors. First, analysis of a publicly-available database (n=1,435) showed that gene expression of specific mitochondrial proteins in ovarian tumors is associated with survival. Quantifying multiple biochemical and molecular markers of mitochondrial content and function in 51 benign and 128 high-grade epithelial ovarian tumors revealed that compared to benign tissue, tumors exhibit 3.3-8.4-fold higher mitochondrial content and respiratory chain enzymatic activities (P<0.001) but similar mitochondrial DNA levels (−3.1%), documenting abnormal mitochondrial phenotypes in tumors. Mitochondrial respiratory chain activity was also associated with interleukin-6 (IL-6) levels in ascites. In benign tissue, negative biobehavioral factors were inversely correlated with mitochondrial content and respiratory chain activities, whereas positive biobehavioral factors tended to be positively correlated with mitochondrial measures, although effect sizes were small to medium (r=-0.43 to 0.47). In contrast, serous tumors showed less pronounced biobehavioral-mitochondrial correlations. These results document abnormal mitochondrial functional phenotypes in ovarian tumors and warrant further research on the link between biobehavioral factors and mitochondria in cancer.

## Introduction

Mitochondria play a pivotal role in tumor growth. Mitochondrial function enables tumor cell adaptation to stressful environments, provides plasticity for tumor growth and survival, confers resistance to apoptotic signaling, and can promote metastatic phenotypes^1–3^. Human tumor cells exhibit altered energy metabolism and metabolic reprogramming associated with alterations in mitochondrial DNA (mtDNA) and its associated proteins^4^. Retrograde signaling from mitochondria to the nucleus can also promote a shift in cellular metabolism towards aerobic glycolysis^5^. Clinically, mitochondrial dysfunction is therefore believed to contribute to tumor progression^6^, chemoresistance^7^, and reduced progression-free survival (PFS)^8,9^. Ovarian tumor cells also secrete pro-inflammatory mediators such as interleukin-6 (IL-6) which in turn promote tumor proliferation, invasion and chemoresistance^10^. Furthermore, metabolic changes in healthy stromal tissue surrounding the tumor (tumor microenvironment) produce metabolites that promote inflammation and support tumor growth^11^. Therefore, mitochondrial function – both within benign and cancerous tissue – represents a potential pathway that may influence tumor behavior and downstream clinical outcomes.

A substantial body of research over the last 30 years documents relationships between biobehavioral factors and cancer progression. Biobehavioral pathways -referring collectively to behavioral, social, and/or psychological factors and concomitant biologic processes -can directly affect signaling mechanisms to enhance tumor growth and impair the immune response in cancer^12^. Specifically, psychological stress, beta-adrenergic and glucocorticoid signaling have been shown to enhance processes such as angiogenesis, tumor cell invasion, and resistance to anoikis, representing additional pathways that promote tumor growth and metastasis^13,14^. Biobehavioral factors also have indirect effects on tumors via effects on host cells in the tumor microenvironment such as macrophages or fibroblasts that can be repurposed to promote tumor growth^12,15,16^. Collectively, these data suggest the existence of biological mechanisms to translate positive and negative psychosocial experiences into molecular changes within tumors.

Mitochondria have been shown to dynamically respond to biobehavioral processes and to be functionally linked to the major neuroendocrine and immune pathways that are known to be modulated by psychological states^17–19^. This includes glucocorticoid stress hormones released during psychosocial stress and their associated metabolic effects^20^. A systematic review of the literature revealed consistent effects of chronic mental stress on mitochondria including mitochondrial content and the activity of respiratory chain complexes involved in energy transformation^21^. In mitochondria, both the respiratory chain involved in energy production and macromolecule biosynthesis, as well as the maintenance of the mitochondrial genome, can be a target of biobehavioral processes. Considering both these genetic and biochemical aspects, we recently developed a high-throughput approach to simultaneously monitor i) mitochondrial content and ii) functional capacity in human blood leukocytes^22^. Using this approach in parallel with daily biobehavioral assessments revealed that mood is associated with mitochondrial functional capacity on subsequent days^22^. Moreover, acute mental stress that induces negative mood can also lead to the release of the immunogenic circulating cell-free mtDNA (ccf-mtDNA) in plasma and serum^23,24^ within minutes, providing further evidence that mitochondria in normal (i.e., non-cancerous) tissues functionally respond to biobehavioral factors.

Here, to examine the potential relevance of mitochondrial function in ovarian cancer, we first surveyed a publicly-available ovarian cancer gene expression dataset and characterized multiple biochemical and molecular markers of mitochondrial content and function in non-cancerous tissue and tissue from ovarian carcinomas, in parallel with inflammation. To explore whether mitochondrial remodeling represents a potential pathway by which biobehavioral processes may support tumor growth and progression, we also quantified the association of biobehavioral factors and mitochondrial phenotypes in tumors or benign tissue.

## Results

### Mitochondrial gene expression and survival

We first leveraged a publicly available gene expression dataset, Kaplan Meier Plotter^25,26^, to examine whether ovarian cancer outcomes, such as survival, are associated with the expression of mitochondrial components. Low levels of a major regulator of mitochondrial metabolism, the mitochondrial pyruvate carrier (MPC2), are known to promote metabolic reprogramming and are associated with greater mortality when MPC is removed in cells and preclinical models^27^. Initial analyses included patients of all histologies, stages, and grades. Among 614 ovarian cancer cases followed over 20 years, we found that relative to those with high MPC2 gene expression, low MPC expression was associated with a 59% increase in mortality (p<0.0001, **Figure 1A**), suggesting that mitochondrial metabolism contributes to ovarian tumor biology in humans.

**Figure 1.**
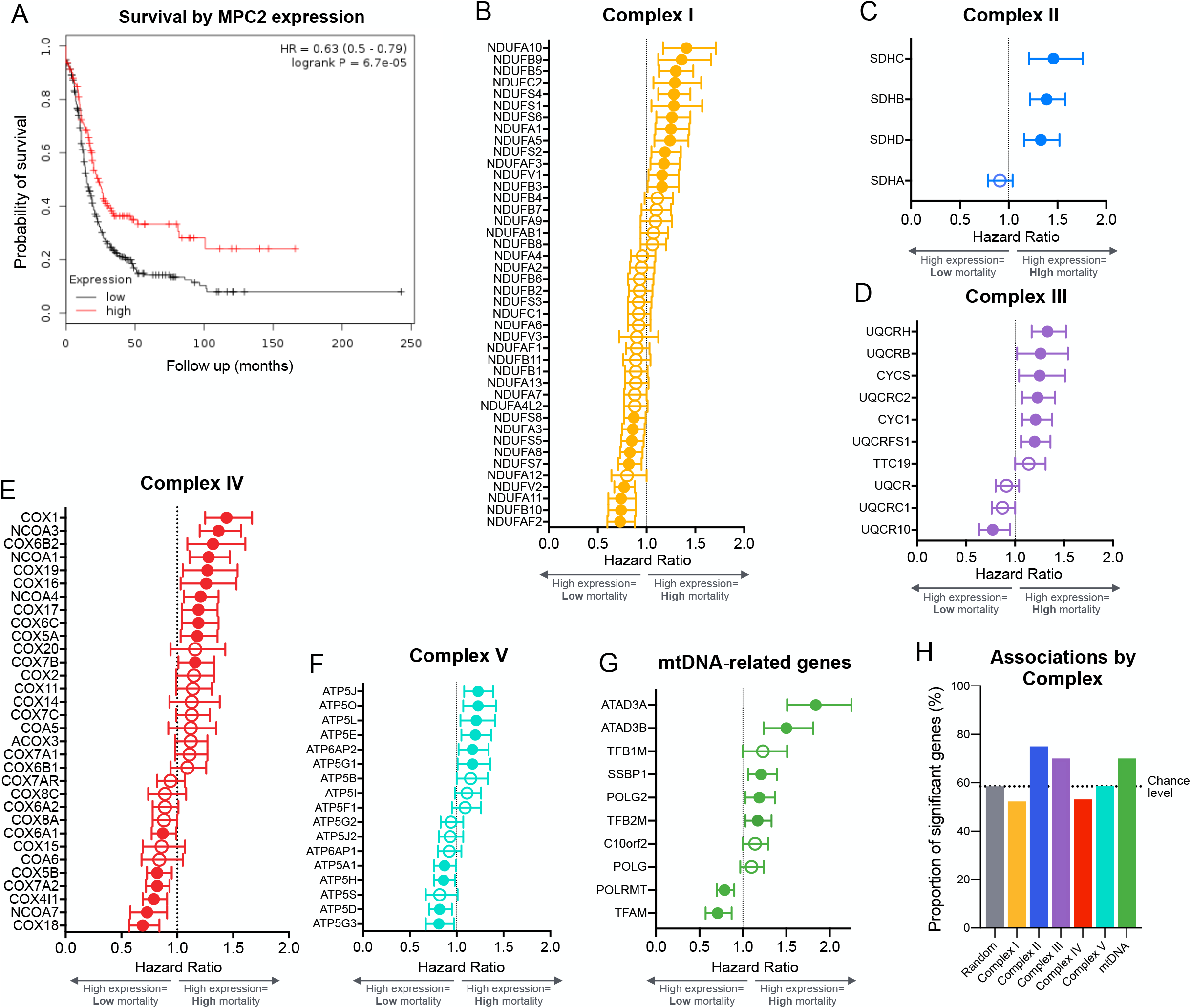
Expression of mitochondrial genes predict survival in ovarian cancer patients. (**A**) Kaplan Meier Plotter (KMPlot.org) survival analysis of patients with ovarian cancer exhibiting low (n=454) or high (n=160) gene expression for a key component of mitochondrial metabolism, the mitochondrial pyruvate carrier 2 (*MPC2*) gene. (**B**) Hazard ratios (HR) from Kaplan Meier survival analysis shown as forest plots for genes encoding subunits of the mitochondrial respiratory chain complexes I, (**C**) II, (**D**) III, (**E**) IV, (**F**) V, and (**G**) and mtDNA-related genes. (**H)** Summary of results illustrating the proportion of significant subunits in previous analysis relative to results from a randomly selected group of 200 genes. Shown are mean HR with 95% C.I. established based on the optimal cutoff; n=614-1435 for each gene.

We then extended this analysis to genes encoding subunits of the respiratory chain complexes I-V, which are involved in mitochondrial oxidative phosphorylation^28^. Similar to MPC2, we generated Kaplan-Meier curves for each available gene encoding a subunit of respiratory chain complexes I through V. The expression levels of respiratory chain subunits, as well as mitochondrial DNA maintenance genes, were variably associated with mortality (**Figure 1B-G**). Compared to a set of randomly selected genes (false discovery rate: 58.5%), the proportion of subunits significantly associated with mortality was higher for complexes II, III and mtDNA-related genes. Sensitivity analyses restricting data to high grade patients performed on expression of MPC2, complex II, and mtDNA-related genes yielded similar results (data not shown). Together, these epidemiological longitudinal data indirectly suggested that ovarian cancer biology in humans may be modulated by metabolic and mitochondrial factors.

### Mitochondrial phenotyping in ovarian tumors

To examine the link between mitochondrial metabolism and ovarian cancer outcomes more directly, we then analyzed tissue from epithelial ovarian tumors (n=128) and benign ovarian masses (n=51) collected during surgery (**Figure 2A**). Ovarian tumors were histologically identified as serous, endometrioid, mucinous, or clearcell, with the majority (78.1%) being serous. All tumors were high grade. Benign masses were of epithelial and stromal histology.

**Figure 2.**
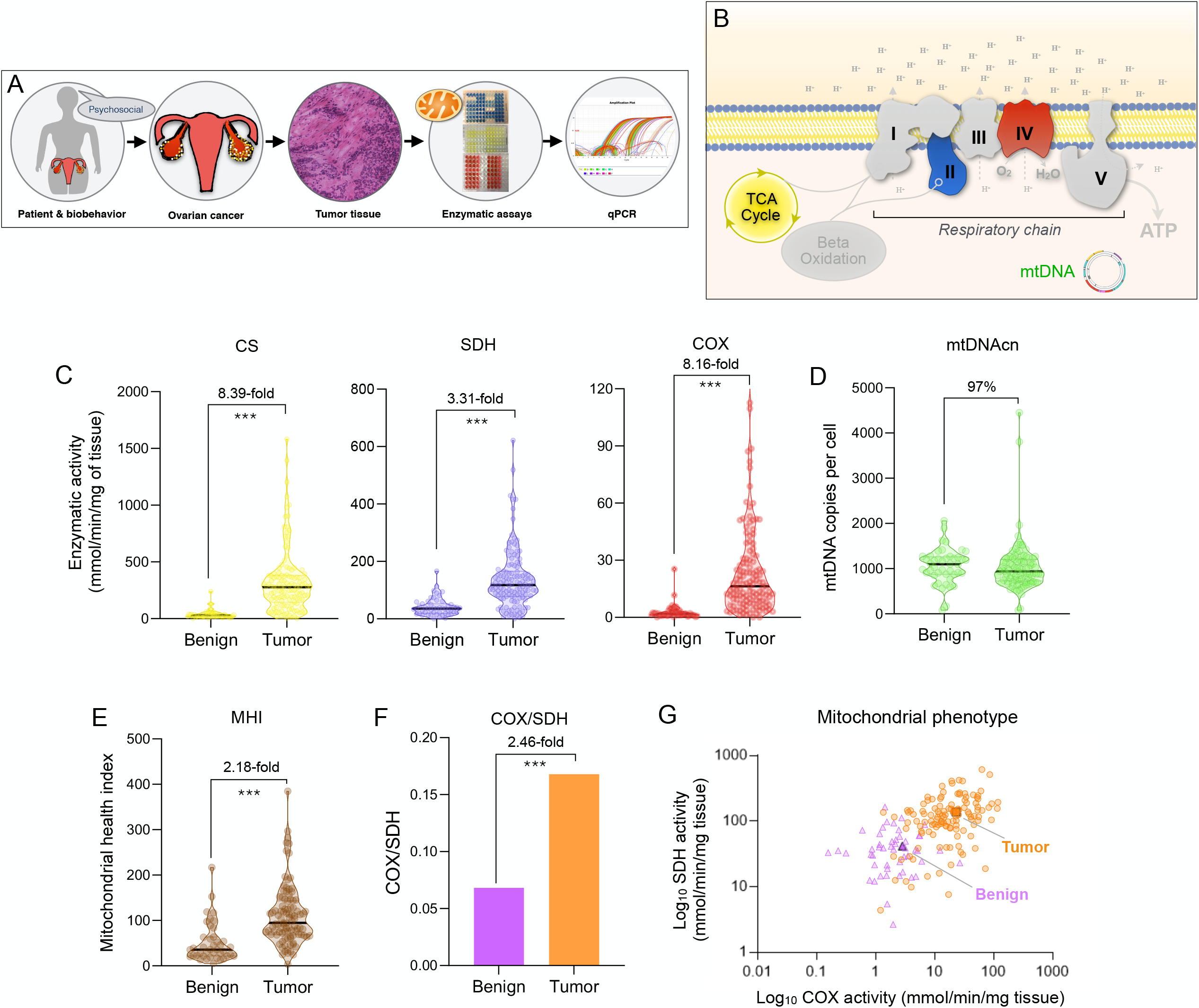
Mitochondrial profiling in benign and tumor tissue. (**A**) Overview of the study design illustrating the multi-level approach ranging from biobehavioral factors to biochemical and molecular assessments of mitochondrial content and function. (**B**) Schematic of the internal components of the mitochondrial respiratory chain (complexes I-V), the tricarboxylic acid (TCA, also Krebs) cycle, and the mitochondrial genome (mtDNA). Enzymatic activities measured include citrate synthase (CS), succinate dehydrogenase (SDH, complex II), and cytochrome *c* oxidase (COX, complex IV). mtDNA copy number (mtDNAcn) was measured using qPCR. Using this data, the Mitochondrial Health Index (MHI) was calculated for each patient with mean-centered activities of respiratory chain complexes, divided by markers of mitochondrial content^3^. (**C**) Enzymatic activities for CS, SDH, and COX, (**D**) mtDNAcn, and (**E**) MHI. (**F**) Average COX/SDH ratio and (**G**) scatterplot illustrating the distinct mitochondrial phenotypes between benign vs tumor. Each data-point represents the average of two measures for a sample. n=51 benign, n=128 high grade tumors, P values from one-way ANOVA performed on log-transformed data. *** p <0.001. The non-transformed data is presented.

To account for possible tumor heterogeneity in the enzymatic and molecular analyses, two partitions of each sample were processed in parallel, measured in triplicate, and the average taken to reflect each ovarian sample most robustly. Our approach included parallel measurements of enzymatic activities for respiratory chain complexes II (succinate dehydrogenase, SDH) and IV (cytochrome *c* oxidase, COX) reflecting mitochondrial respiratory chain capacity, as well as citrate synthase (CS) and mtDNA copy number (mtDNAcn) reflecting mitochondrial content (**Figure 2B**). On average, in tumor the mitochondrial enzymes were moderately correlated with each other, sharing 20.3-47.3% of their variance, whereas mtDNAcn was minimally correlated with enzymatic activities (1.1-3.4% of shared variance). These data suggest that each mitochondrial marker contributes different information about the tumor mitochondrial phenotypes and that mtDNAcn is likely regulated independently from other measures. Age was examined as a possible factor that might influence mitochondrial function. Enzymatic activities and mtDNAcn in tumor were minimally associated with age (r^2^ = 0.00-0.078), such that <8% of the variation in mitochondrial content and function was attributable to age in this cohort. The results were similar for associations of enzymatic activities and mtDNAcn with age in benign tissue (r^2^ = 0.00-0.043).

### Mitochondrial content and function are elevated in tumor relative to benign

We first compared mitochondrial measures between high grade ovarian tumor tissue and benign ovarian masses. Relative to benign tissue, ovarian tumors exhibited a striking 8.4-fold higher CS activity, 3.3-fold higher SDH activity, and a 8.2-fold higher COX activity (**Figure 2C**), but a similar (3.1% lower, N.S.) mtDNAcn (**Figure 2D**). The mitochondrial health index (MHI), a computed variable representing the ratio of respiratory chain activity (SDH and COX) divided by markers of mitochondrial content (CS and mtDNAcn)^22^, was also double in tumors compared to benign tissue (**Figure 2E**). Because SDH is uniquely encoded in the nuclear genome, whereas COX is partially mtDNA-encoded, we also computed the ratio of both enzymes (**Figure 2F**), which can be indicative of abnormal mitochondrial phenotypes^29^. Here the COX/SDH ratio was approximately 3 times higher in tumors, and plotting each tumor and benign sample in a bi-plot showed fairly good separation between both groups, indicating a possible imbalance or recalibration of the respiratory chain components in tumor tissue (**Figure 2G)**. Moreover, the minimal decline in mtDNAcn in light of the approximately 3.3-8.4-fold higher mitochondrial mass means that the mitochondrial genome *density* (mtDNA/CS) per mitochondrion is dramatically reduced in tumors. Together, these findings demonstrate that beyond gross changes in mitochondrial abundance, tumor mitochondria exhibit abnormal mitochondrial phenotypes that are qualitatively different from benign tissue.

### Mitochondrial content and function across tumor stages and types

Overall, tumors presented with a broad range of mitochondrial enzymatic activities and mtDNAcn (**Figure 3A-B**). Average respiratory chain enzyme activities were approximately equivalent in Stages I through III and then decreased substantially by Stage IV. For CS, SDH, and COX, metastatic tumors (Stage IV) showed approximately half (55.8-61.9%) the enzymatic levels of stage I tumors, although these differences were non-significant (*p* values > 0.05) due to high variation within each stage. mtDNAcn of Stage IV tumors was on average 28.2% lower than in Stage I tumors, or half of the magnitude of differences between stages I and IV observed for enzymatic activities) (**Figure 3C**), again suggesting that mtDNAcn may be regulated independently from enzymatic components. The MHI did not differ statistically across stages (**Figure 3D**).

**Figure 3.**
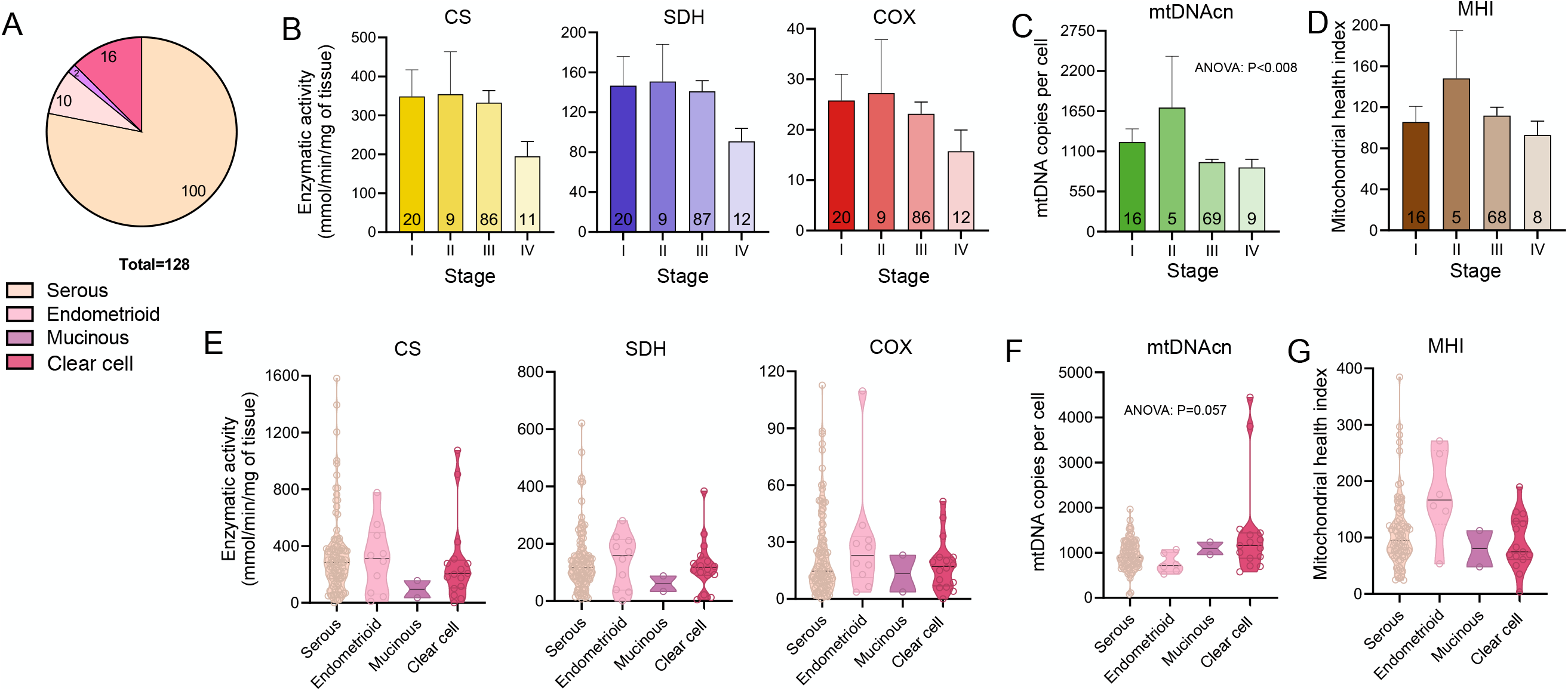
Mitochondrial phenotyping across tumor types and benign tissue. (**A**) Tumor types assayed in this study. (**B**) Mitochondrial enzymatic activities, (**C**) mtDNAcn, and (**D**) MHI by tumor stage. Data are means ± SEM. (**E**) Enzymatic activities, (**F**) mtDNAcn, and (**G**) MHI values by tumor type shown as violin plots with each datapoint representing a tumor or individual patient. P values from one-way ANOVA performed on log-transformed data. The non-transformed data is presented.

Next, we compared mitochondrial measures across tumor histologies. This revealed two main points: i) there is large inter-individual variation in enzymatic activities even within the same tumor type; and ii) there are potentially large differences in activities between tumor histologies (**Figure 3E-G**), although this study was not powered to detect these differences.

### Tumor mitochondrial parameters and IL-6

Mitochondrial function is linked to pro-inflammatory signaling and cytokine production, including interleukin 6 (IL-6)^30,31^ and IL-6 signaling may influence tumor-promoting processes such as angiogenesis^32^. Therefore, to evaluate whether mitochondrial measures are related to the pro-inflammatory phenotype, we measured IL-6 levels in ascites and plasma from participants with ovarian tumors, across all stages, and systematically tested their association with tumor mitochondrial respiratory chain enzymatic activities and mtDNAcn (**Figure 4A**). Tumor mitochondrial respiratory chain activity and content, particularly the integrated metric MHI (**Figure 4B**), were consistently negatively correlated with IL-6 levels in ascites (*p* values = 0.042 to < 0.001); moreover correlations were up to an order of magnitude larger in ascites than in plasma. In contrast, mtDNA density per mitochondrial unit (ratio of mtDNAcn/CS) was correlated positively with ascites IL-6 (*p* value = 0.014) **(Figure 4C)**, possibly reflecting the pro-inflammatory signaling effects of mtDNA^33,34^.

**Figure 4.**
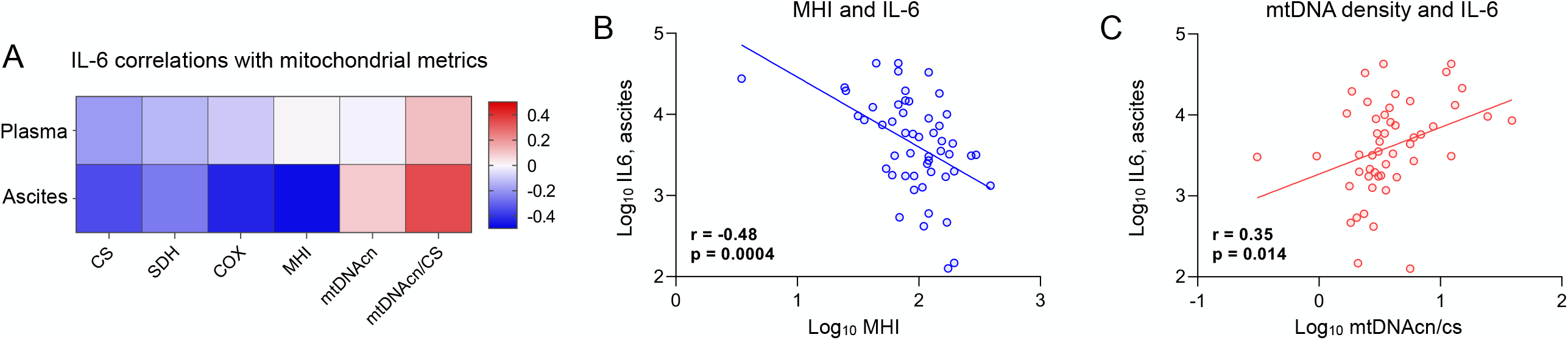
Mitochondrial measures and IL-6 in high grade tumors. (**A**) Pearson correlations between mitochondrial measures and IL-6 levels measured in ascites and plasma. Note the similar patterns of correlation between ascites and plasma, and the contrast in the associations between mtDNA-related measures (positive) and primary measures of mitochondrial content and respiratory chain function (negative). (**B**) Scatterplots of the correlations between ascites IL-6 levels and the mitochondrial health index (MHI), and (**C**) mtDNA density per mitochondrion (mtDNAcn/CS). n=50-64 for ascites, 82-110 for plasma.

### Associations between biobehavioral factors and mitochondrial phenotypes

Because serous tumors are the predominant tumor type in ovarian cancer, we focused analyses with biobehavioral factors on high grade serous tumors. These exploratory analyses examined whether mitochondrial parameters were associated with negative and positive biobehavioral factors. Negative biobehavioral factors were moderately inter-correlated, sharing between 14.9 and 51.9% of their variance, and the proportion of shared variance among positive biobehavioral factors ranged from 0.003 to 42.1%, indicating no co-linearity to moderate co-linearity. The strength and direction of each mitochondria-biobehavioral correlation is displayed as a heat map in **Figure 5A** and with 95% confidence intervals shown for the enzymatic mitochondrial measures (CS, SDH, COX) in **Figure 5B**. These results suggest patterns of association with relatively small to moderate effect sizes (range: r=-0.43 to 0.47) across multiple mitochondrial measures and biobehavioral assessments, leading to three main observations.

**Figure 5.**
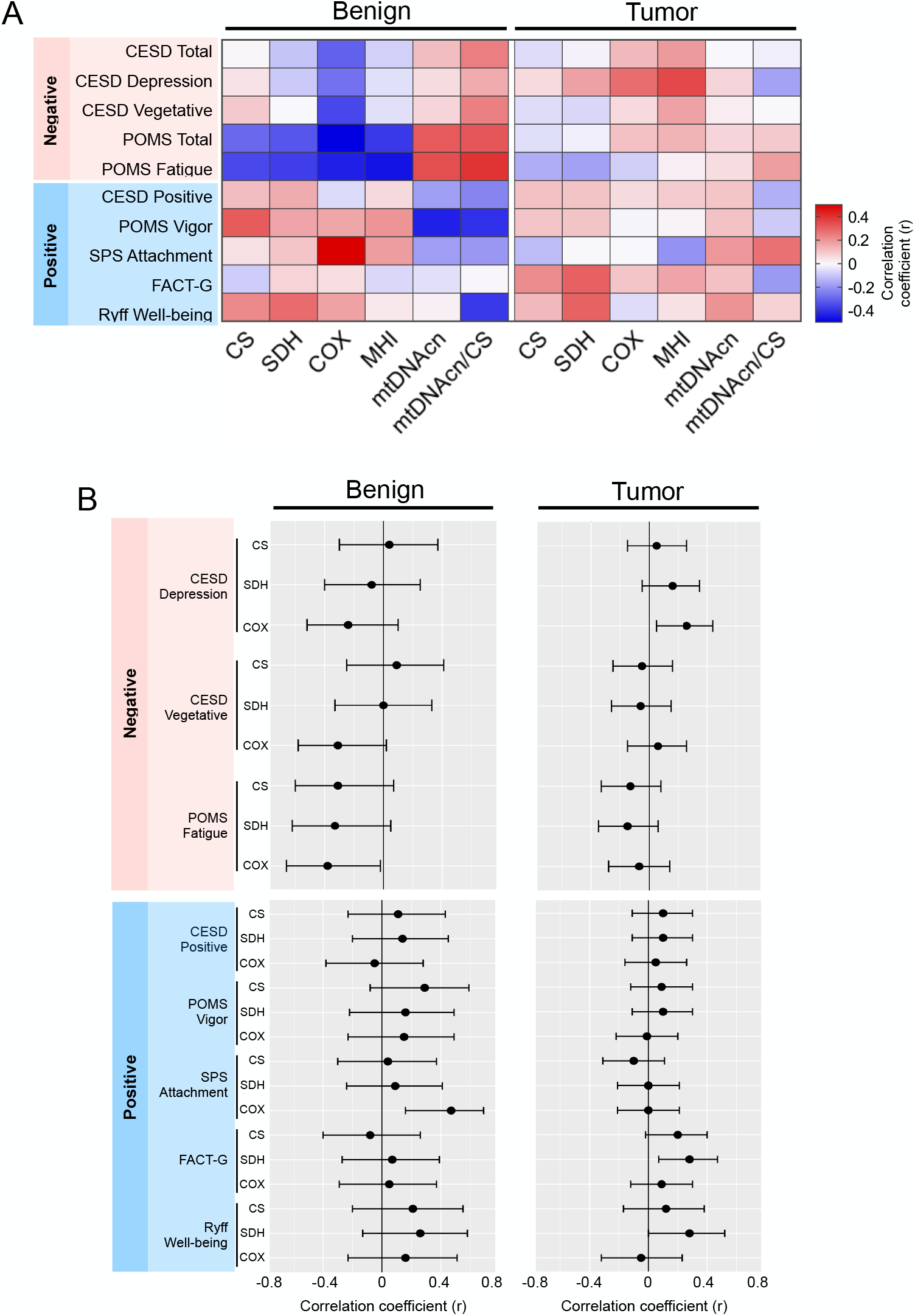
Associations of biobehavioral factors with mitochondrial content and function in high grade serous tumor and in epithelial and stromal benign tissues. (**A**) Correlations between negative/positive biobehavioral factors and markers of mitochondrial content (CS, mtDNAcn), mitochondrial respiratory chain enzymatic activities (SDH, COX), and mtDNAcn density (mtDNAcn/CS). Results are Pearson correlation coefficients illustrated in a heatmap depicting the direction (color) and strength (saturation) of the associations. Benign and high grade serous tumor tissues may have opposite associations among biobehavioral factors and mitochondrial activity particularly for CS and respiratory chain complexes II (SDH) and IV (COX). (**B**) Effect sizes shown in A for the primary enzymatic measures (CS, SDH, COX) with 95% confidence intervals. Whereas data from subscales and scales are presented in A, overlapping scales and subscales are omitted from this panel. Correlation coefficients determined from log-transformed data. n=22-36 benign, n=37-91 tumors; correlations are Pearson r from log-transformed values.

First, in benign tissues, negative biobehavioral measures such as depression, fatigue, and negative mood tended to be generally associated with lower levels of mitochondrial enzymatic activities. Fatigue was the measure most consistently related to mitochondrial activity, with effect sizes in the moderate range for all mitochondrial enzymes, and significant negative associations with COX *r*=-.38, *p*=0.04 and MHI *r*=-0.40, *p*=0.039 and positive associations with mtDNA/CS *r*=.37, *p*=0.059. In contrast, positive biobehavioral measures such as vigor, well-being, and positive mood tended to be positively correlated with mitochondrial parameters, although only the relationship between social attachment and COX reached statistical significance, *r*=0.47, *p*=0.004. Although these relationships were not generally statistically significant due to the small sample size (n=27-36) in many of these analyses, and effect sizes are small to moderate, we note an apparent consistency in their direction across primary mitochondrial measures (CS, SDH, COX). This would seem to be consistent with previous findings in blood leukocytes^22^, and these observations should be examined further in more strongly powered studies.

Second, in benign tissue, compared to the associations seen in respiratory chain enzymes, associations between mtDNAcn and biobehavioral factors appear to be in directions opposite to those noted above. This would be consistent with the existence of independent mechanisms linking biobehavioral factors to mtDNAcn and respiratory chain function.

Third, in contrast to the apparent biobehavioral-mitochondrial associations in benign tissue, these associations in high grade serous tumor tissue tended to be smaller in magnitude (see Figure 5B). In tumor, contrary to expectation, both positive and negative factors were associated with higher levels of respiratory chain enzymes (depressive mood and COX: *r*=0.26, *p*=0.014; FACT quality of life (QOL) and SDH (*r*=.29, *p*=0.009); well-being and SDH (*r*=.28, *p*=0.050). Interestingly, biobehavioral characteristics indicating positive affect (CESD-positive, POMS-vigor), which may be more fleeting, were not strongly associated with respiratory chain enzymes. In contrast, the positive dimensions of QOL and well-being which may reflect a more sustained positive orientation, were significantly associated with higher levels of respiratory chain activity in tumor tissue.

## Discussion

In this study of epithelial ovarian cancer, we examined various enzymatic and molecular mitochondrial measures and the intersection of mitochondrial biology and biobehavioral factors. After interrogating mitochondrial gene expression from a large database of ovarian cancer survival as proof-of-concept of the significance of mitochondria in ovarian tumor biology, we leveraged a high-throughput mitochondrial phenotyping platform to measure multiple markers of mitochondrial content and function. In parallel, to investigate whether mitochondrial remodeling represents a potential pathway by which biobehavioral processes may support tumor growth and progression, we analyzed several domains of psychosocial functioning in ovarian cancer patients at the time of surgery. From a tumor biology perspective, our mitochondrial respiratory chain and mtDNA results demonstrate striking differences in mitochondrial phenotypes between tumor and benign tissues, and suggest potential differences among different histologies and tumor stages. Our exploratory analyses also suggest relatively blunted associations between biobehavioral factors and mitochondria in high grade serous tumors, in comparison with benign tissue where positive and negative biobehavioral measures are potentially more strongly related to mitochondrial function. This is the first study to examine the association of biobehavioral factors with mitochondrial content and function in cancer. Overall, these findings extend pre-clinical work suggesting mitochondrial reprogramming in ovarian cancer progression in humans.

The extension of pre-clinical findings concerning the influence of MPC2 on mitochondrial metabolism and tumor growth^27^ to MPC2 gene expression and survival in women with ovarian cancer provided proof-of-concept that mitochondrial (dys)function may contribute to clinical outcomes. In systematically broadening the search to other mitochondrial components, one of the striking observations from the gene expression-survival analysis (Figure 1) is the heterogeneity in the direction of the associations for proteins belonging to the same complex. To our knowledge, this has not previously been examined in detail. For example, higher expression of NDUFAF2 was associated with lower mortality. NDUFAF2 is a complex I assembly factor that is required for synthesizing the part of complex I that protrudes in the mitochondrial matrix and binds NADH (N module) produced by the Krebs cycle. Paradoxically, we find that an elevated expression of many of the *structural* complex I subunits that are part of the N module is associated with higher mortality. This may be due to the fact that when such subunits are overexpressed in the absence of a concomitant overexpression of NDUFAF2, they interact with other proteins to disrupt other signaling networks. For instance, NDUFV1 interacts with LRRK2^35^, and NDUFS1 interacts with the iron-sulfur cluster co-chaperone protein HscB^36^; hence it is conceivable that when overexpressed they bind and sequester HscB to the point that other cellular processes dependent on iron-sulfur clusters are compromised. Similarly, NDUFS6 interacts with BOLA3, which also regulates iron-sulfur cluster biogenesis^37^ while NDUFS4 interacts with proline-rich transmembrane protein 2 (PRRT2)^38^. In relation to mtDNA, higher expression of both isoforms of ATAD3 (ATAD3A and ATAD3B), which are likely involved in cholesterol metabolism in addition to mtDNA nucleoid regulation^39^, is associated with >50% increase in mortality. In contrast, expression of both TFAM and the mitochondrial RNA polymerase, which both promote mtDNA gene expression^40^, are negatively associated with mortality. Thus, as expected from known protein-protein interactions, these data also demonstrate that different respiratory chain subunits belonging to the same protein complexes could have vastly different influence on tumor biology depending upon their specific molecular interactions and position within multi-protein respiratory chain complexes.

One point of interest relates to mitochondrial content, here inferred from the combination of multiple markers^41^, found to be exceptionally high in ovarian cancers relative to benign tissue. While some studies have reported a decrease in SDH and COX activity in cancers like breast cancer and renal carcinoma^42^, we report increased activity in ovarian cancer compared to healthy tissue. CS upregulation has been observed in other ovarian cancer studies^41^ In cancer cells, mitochondria are biosynthetic centers -several aspects of their function, including the respiratory chain and associated Krebs cycle enzymes are diverted to provide biosynthetic products to support cell growth^6,43^ (reviewed in^44^). The upregulation of CS (citrate synthase) might be responsible for regulating citrate levels and affect cellular lipid synthesis, which drives key cellular processes such as transformation, tumor development, and disease progression^45^. Citrate, like other mitochondria-derived metabolites with pro-oncogenic effects^46^, is considered an oncometabolite^42^. Other known mitochondrial metabolites accumulated in tumors can also activate oncogenic signaling cascades, including succinate and fumarate that are the substrate and product, respectively, of the reaction catalyzed by SDH (succinate dehydrogenase)^47^,^46^. As increased CS and SDH activity would result in an increase in citrate and fumarate, our findings indicating a significant increase in CS and SDH activities in tumor compared to benign mitochondria suggest that this may be worth examining in future studies. More generally, increased mitochondrial mass could be the product of increased biogenesis, decreased autophagic removal of mitochondria, or a combination of these and other factors. In support of this point, mitochondrial biogenesis has been observed in various human tumor types and correlates with metastatic behavior^48^ and poor clinical outcomes^2^.

Another unresolved point relates to the association between mitochondrial content and function with IL-6. Besides abnormal mitochondrial metabolism, inflammation is another biological hallmark of tumor transformation and progression^49^. In cancer patients, we noted a negative association between mitochondrial enzyme activities and IL-6 levels in ascites. In circulating blood, these associations were in the same direction as ascites but less robust, consistent with the tumors being the primary source of IL-6^50^ and the “leakage” of IL-6 into circulation. Our data suggest a fairly robust negative association between both mitochondrial mass and respiratory chain function and IL-6. This apparent anti-inflammatory phenotype of mitochondria-rich tissues could be explained by a number of mechanisms that require further work to confirm. One possibility is that this association reflects some crosstalk between tumor mitochondria and resident immune cells^51^,52. Consistent with our data, the upregulation of mitochondrial biogenesis (higher mitochondrial mass, respiratory chain activity) promotes an anti-inflammatory shift in monocytes^53^. Acute perturbations of mitochondrial respiratory chain function also alter cytokine production in human leukocytes^54^, substantiating the link between mitochondrial function and cytokine production in human immune cells. Together, these data suggest that the mitochondrial phenotype may influence the inflammatory state, rather than the other way around. However, further work is needed to test this proposition in ovarian cancer.

There is a need to establish biologically plausible pathways linking psychosocial experiences with human health outcomes^55^. Here, the exploratory associations between biobehavioral and mitochondrial measures suggest that compared to benign tissue, ovarian tumors may respond differently to negative psychosocial factors. Benign tissue appeared to have similar associations to those we have previously reported in leukocytes in response to positive and negative biobehavioral factors^22^. Specifically, positive mood (more so than negative mood) was significantly associated with future mitochondrial respiratory chain function (i.e., MHI). While the basis for any such difference remains unclear, we speculate that abnormal mitochondrial phenotypes in tumors may render tumor mitochondria less responsive to the natural influence of neuroendocrine factors believed to mediate the effects of psychosocial factors in other cell types^12^. However, on the basis of small effect sizes that are largely not statistically significant, we hesitate to draw definite conclusions, and we advocate for further, more adequately-powered studies to examine this question.

Contrary to observations involving mitochondrial content and respiratory chain activity, mitochondrial genome abundance (mtDNAcn), particularly when expressed as mtDNA density per mitochondrion (relative to mitochondrial mass, mtDNAcn/CS), was found to be positively correlated with ascites IL-6 levels. This antagonistic association of mitochondrial respiratory chain components and mtDNAcn with IL-6 is similar to our findings with biobehavioral factors. Interestingly, mtDNA instability can cause its release in the cytoplasm and extracellular space where it is potentially recognized as foreign (i.e., bacterial) by innate immune response systems, triggering the production of antiviral and pro-inflammatory cytokines^56,57^. Although it was not possible to measure levels of cell-free mtDNA in the current study, our data suggest that mtDNA signaling could, in ovarian cancer, have pro-inflammatory effects. The origin of this opposite association between respiratory chain components and mtDNAcn with both inflammation and biobehavioral outcomes remains unclear but emphasizes the notion that respiratory chain function and mtDNAcn are not equivalent markers of mitochondrial function.

Overall, our results demonstrate abnormal mitochondrial phenotypes in ovarian tumors and open new questions related to the role of mitochondrial biology in ovarian cancer, including their sensitivity to biobehavioral factors. As mitochondrial dysfunction is directly implicated in several chemoresistance pathways^7^, future work may also examine how altered mitochondrial functioning may contribute to clinically relevant processes such as chemoresistance and clinical outcomes.

## Methods

### Publicly available ovarian cancer gene expression dataset

To examine the association between tumor gene expression and progression-free survival, we queried Kaplan Meier Plotter (http://kmplot.com) to compute the survival over time in 1,435 ovarian cancer patients. Kaplan Meier Plotter is the largest publicly available dataset of microarray-based gene expression in 21 different cancer types, including ovarian cancer. Analyses were not restricted to different subtypes (histology, stage, grade, TP53 mutation) or to treatment groups (debulk, chemotherapy). We analyzed the prognostic value of transcript level for genes encoding mitochondrial respiratory chain subunits of complexes I through V, as well as genes related to mtDNA maintenance and stability. Patients were divided into two groups of high and low gene expression using default settings to establish the optimal threshold as a cutoff. The two groups were compared on progression-free survival over follow up periods of up to 20.8 years. To obtain an unambiguous expression estimate for each gene, the optimal probe selected based on the JetSet algorithm^58^ was selected to represent each gene. A survival plot was then generated for each gene with sample sizes varying according to the gene data available (n=614-1435 for each gene). The KM survival estimates and log-rank tests statistic were used to determine the hazard ratio, 95% CI and p-value for each gene, and the data plotted as forest plots to visualize the direction and magnitude of associations between complex-specific genes and progression-free survival. To obtain an estimate of false discovery rate in this dataset, we randomly selected 200 genes (“null set”) out of the total 15,000 available human genes. We obtained the hazard ratios and 95% confidence interval for each random gene and computed the proportion of significant genes with these parameters, which serves as a reference to evaluate the proportion of significant genes in each mitochondrial respiratory chain complexes and mtDNA-related genes. Sensitivity analyses restricting the data to high grade patients (KMPlotter grades 2 and 3) were then conducted on MPC2 and Complex II- and mtDNA-associated genes which showed > 70% significant associations with survival compared to the null set.

### Clinical Data

Participants were part of a larger study of biobehavioral factors and tumor progression in ovarian cancer who were prospectively recruited at an initial clinic visit pre-surgery/treatment initiation at 2 large Midwestern hospitals between 12/03 and 8/14. All procedures were IRB approved and all participants signed informed consents. All human subjects were approved by the Institutional Review Boards at the University of Iowa and Washington University and all research was performed in accordance with relevant guidelines and regulations.

#### Inclusion/exclusion

Participants over 18 years old with a newly-diagnosed, histologically-confirmed, primary invasive epithelial ovarian, peritoneal, or fallopian tube carcinoma or a tumor suspected for ovarian cancer which turned out to be benign were eligible for inclusion in the parent study. Primary cancers of another organ, non-epithelial ovarian tumors, tumors of low malignant potential, previous cancer, use of systemic steroid medication in the last 4 months, or comorbidities known to influence the immune response were excluded. For the current analyses, only high grade ovarian tumor tissue was included, and benign tissue was restricted to epithelial (e.g. cystadenoma) and stromal (e.g. fibroma) histology.

#### Procedures

Psychosocial assessments were completed at home prior to surgery or initiation of treatment and clinical data was obtained from medical records. Tumor samples were obtained from primary surgeries conducted prior to beginning of treatment. Tumor samples were flash frozen in liquid nitrogen immediately upon acquisition. Tissue was available from 128 patients subsequently diagnosed with high grade epithelial ovarian cancer and from 51 patients with benign epithelial or stromal pelvic masses. Plasma was obtained immediately prior to surgery and ascites was obtained from surgery. Both were frozen at −80°C until analysis.

### Psychosocial measures

The Center for Epidemiological Studies-Depression Scale (CES-D) contains 20 items assessing frequency of depressive symptoms over the past week^59,60^. Scores of 16 or higher are indicative of clinical depression. Four validated subscales have been described and associated with physiological variables in previous ovarian cancer research^59,60^. In this study, we used the total score, and the depressed mood, positive mood, and vegetative subscales.

The Profile of Mood States short form (POMS-SF) includes 37 adjectives describing mood over the last month. Six factors are included in this measure: anxiety, dysphoria, anger, vigor, fatigue, and confusion. A total distress score (total mood disturbance: TMD) is derived from the sum of the 5 negative mood factors minus the vigor subscale. The POMS-SF is well-validated, has strong psychometric properties, and is commonly used for assessment of distress in cancer populations^61^. Here, in addition to the total scale, we used the fatigue and vigor scales as ways of examining energy/vitality levels that might be associated with mitochondrial activity.

The Social Provisions Scale (SPS) is a 24-item self-report scale measuring an individual’s perceptions of their social support^62^. Here we use the attachment subscale which has shown strong associations with physiological measures in previous studies of female cancer patients^63^.

The Psychological Well-Being Scale (PWBS) measures perceived psychological well-being^64^. We administered 4 of the 6 subscales ^(^purpose in life, environmental mastery, personal growth, and self-acceptance) thought to be particularly relevant for individuals with chronic health challenges. We used the 7-item version of the scales, with higher scores indicating greater well-being.

The Functional Assessment of Cancer Therapy (FACT-G) is a well-validated and psychometrically sound 27-item scale assessing 4 facets of quality of life (physical, emotional, functional, and social well-being) experienced by cancer patients during the previous week. Higher scores indicate better quality of life^65^.

### Mitochondrial content and function

#### Tissue preparation

Tumor samples were flash frozen and stored in liquid nitrogen until processing. For each tumor or benign sample, two partitions of 50-65 mg were cut with a scalpel on a clean, cooled surface. The partitions were allowed to thaw and were mechanically minced on ice. Minced samples were placed on dry ice and stored at −80°C until homogenization in batches. To account for potential tumor heterogeneity, enzymatic activities and mtDNAcn were measures on each partition individually and the average of both partitions was calculated, producing a single, more stable value for each tumor and benign sample. Pre-cut tumor samples were mechanically homogenized with Tungsten beads (Cat #69997) in homogenization buffer containing 1 mM EDTA and 50 mM triethanolamine (pH 7.4) 1:20 (1mg/20ul), using a robotized TissueLyser II (Qiagen) with a 1-minute run (30 cycles/sec) followed by an incubation on ice for 5 min, repeated for a total of three runs. Samples were vortexed before each assay to ensure homogeneity.

#### Mitochondrial enzymatic activity assays

Mitochondria enzyme activities were quantified spectrophotometrically for citrate synthase (CS), cytochrome c oxidase (COX, complex IV), succinate dehydrogenase (SDH, complex II), and NADH dehydrogenase (complex I). For each assay, 20 ul of homogenate was added per reaction. All assays were performed at 30°C. *Citrate synthase* (CS) activity was measured by detecting the increase in absorbance at 412 nm, in a reaction buffer (200 mM Tris, pH 7.4) containing acetyl-CoA 0.2 mM, 0.2 mM 5,5’-dithiobis-(2-nitrobenzoic acid) (DTNB), 0.55 mM oxaloacetic acid, and 0.1% Triton X-100. Final CS activity was obtained by integrating OD^412^ change from 120-300 sec, and by subtracting the non-specific activity measured in the absence of oxaloacetate. *Cytochrome c oxidase (COX, or complex IV)* activity was measured by detecting the decrease in absorbance at 550 nm at 30°C, in a 100mM potassium phosphate reaction buffer (pH 7.5) containing 0.1% n-dodecylmaltoside and 100uM of purified reduced cytochrome c. Final COX activity was obtained by integrating OD^550^ change over 100-350 sec and by subtracting spontaneous cyt c oxidation without cell lysate. *Succinate dehydrogenase (SDH, or complex II)* activity was measured by detecting the decrease in absorbance at 600 nm at 30°C, in potassium phosphate 100 mM reaction buffer (pH 7.5) containing 2 mM EDTA, 1mg/ml bovine serum albumin (BSA), 4 μM rotenone, 10 mM succinate, 0.25 mM potassium cyanide, 100 μM decylubiquinone, 100 μM DCIP, 200 μM ATP, 0.4 μM antimycin A. Final SDH activity was obtained by integrating OD^600^ change over 200-600 sec and by subtracting activity detected in the presence of malonate (5 mM), a specific inhibitor of SDH. *NADH dehydrogenase (complex I)* activity was measured by detecting the decrease in absorbance at 600 nm at 37°C, in potassium phosphate 100 mM reaction buffer (pH 7.5) containing 2 mM EDTA, 3.5 mg/ml bovine serum albumin (BSA), 0.25 mM potassium cyanide, 100 μM decylubiquinone, 100 μM DCIP, 200 μM NADH, 0.4 μM antimycin A. Final complex I activity was obtained by integrating OD^600^ change over 200-440 s and by subtracting activity detected in the presence of rotenone (500 μM) and piericidin A (200 μM), specific inhibitors of complex I. Activities for complex I were too low and were not used for analyses. The molar extinction coefficients used were 13.6 L mol^-1^cm^-1^ for DTNB, 29.5 L mol^-1^ cm^-1^ for reduced cytochrome c, and 16.3 L mol^-1^ cm^-1^ for DCIP. Samples were lysed for 10 hours at 55°C, followed by inactivation at 95°C for 10 minutes. The lysate was directly used as template DNA for measurements of mtDNA copy number.

#### mtDNA copy number

The same homogenate used for enzymatic measurements (10ul) was lysed in lysis buffer containing 100mM Tris HCl pH 8.5, 0.5% Tween 20, and 200ug/ml proteinase K (10 hours at 55°C, and used directly for qPCR-based measurements of mtDNA copy number. qPCR reactions were setup in triplicates using a liquid handling station (ep-*Motion*5073, Eppendorf) in 384 well qPCR plates. Duplex qPCR reactions with Taqman chemistry were used to simultaneously quantify mitochondrial and nuclear amplicons in the same reactions. Master Mix_1_ for ND1 (mtDNA) and B2M (nDNA) included: TaqMan Universal Master mix fast (life technologies #4444964), 300nM of custom design primers and 100nM probes (ND1-Fwd: GAGCGATGGTGAGAGCTAAGGT, ND1-Rev:CCCTAAAACCCGCCACATCT,Probe: HEX-CCATCACCCTCTACATCACCGCCC-3IABkFQ.B2M-Fwd: CCAGCAGAGAATGGAAAGTCAA, B2M-Rev: TCTCTCTCCATTCTTCAGTAAGTCAACT, Probe: FAM-ATGTGTCTGGGTTTCATCCATCCGACA-3IABkFQ). Master Mix_2_ for COX1 (mtDNA) and RNaseP (nDNA) included : TaqMan Universal Master Mix fast, 300nM of custom design primers and 100nM probes (COX1-Fwd: CTAGCAGGTGTCTCCTCTATCT, COX1-Rev: GAGAAGTAGGACTGCTGTGATTAG, Probe: FAM-TGCCATAACCCAATACCAAACGCC-3IABkFQ. RnaseP-Fwd: AGATTTGGACCTGCGAGCG, RNaseP-Rev: GAGCGGCTGTCTCCACAAGT, Probe: FAM-TTCTGACCTGAAGGCTCTGCGCG-3IABkFQ. The samples were then cycled in a QuantStudio 7 flex qPCR instrument (Applied Biosystems) at 50°C for 2min, 95°C for 20sec, 95°C for 1min, 60°C for 20sec for 40x cycles. Reaction volumes were 20ul. To ensure comparable Ct values across plates and assays, thresholds for fluorescence detection for both mitochondrial and nuclear amplicons were set to 0.08.

mtDNAcn was calculated using the ΔCt method. The ΔCt was obtained by subtracting the average mtDNA Ct values from the average nDNA Ct values. Relative mitochondrial DNA copies are calculated by raising 2 to the power of the ΔCt and then multiplying by 2 to account for the diploid nature of the nuclear genome. The number of mtDNAcn = 2^ΔCt^ x 2. Both ND1 and COX1 yielded highly correlated results and the average of both amplicon pairs was used as mtDNAcn value for each sample.

### Statistical analyses

The hazard ratios (HR) and 95% confidence for survival differences between high- and low-expression tumors was obtained from http://kmplot.com. Statistical Package for the Social Sciences (SPSS) version 25.0 and R version 3.6.3 were used to analyze all other data. All distributions were examined for outliers and nonnormality. Log transformations were applied to normalize distributions where necessary. Mitochondrial markers were compared between benign and tumor data using analyses of variance (ANOVAs). ANOVAs were also used to examine overall differences in means of mitochondrial variables according to cancer stage or histology. Pearson product moment correlations were calculated to assess relationships between psychosocial variables and mitochondrial variables and between mitochondrial variables and IL-6. A 95% confidence interval was generated for each correlation coefficient.

## Author contributions

S.L. designed and obtained funding for the parent study. M.P. designed the mitochondrial phenotyping analyses. P.H.T. provided project administration and clinical data. M.G. provided resources and clinical data. L.D. provided pathology review. S.B., M.A.M., M.K.T., and A.T. performed mitochondrial phenotyping. S.L., S.B., M.P. and R.T.O. performed analyses. S.B., M.P., and S.L. prepared figures. M.P., S.L. and S.B. drafted manuscript. E.O.A. and A.K.S. contributed to interpretation of results. All authors reviewed the manuscript.

## Funding and acknowledgements

This work was supported in part by Grants CA193249 and CA193249S1 (S.L.) and 3P30CA086862 (George Weiner) from the National Cancer Institute, and by the Wharton Fund and NIH grant GM119793 (M.P.).

